# Unveiling the secrets of brain resilience: Multimodal neuroimaging insights into Neural dynamics and neuroprotection in ageing

**DOI:** 10.64898/2025.12.21.695787

**Authors:** Bhaskar Roy, Prasun K. Roy

## Abstract

With the onset of ageing the likelihood of brain degeneration increases but so does the remarkable capability of the brain to reorganize itself termed as neurocognitive resilience. Leveraging the neuroimaging mapped anatomical signatures to demarcate age related alteration of an individual brain is a promising strategy to identify ageing phenotypes and estimate the molecular cues account for coping mechanisms.

This study endeavour to identifying neuroprotective signatures using multimodal structural and functional brain scans of healthy aged male and female cohort across dorsal-ventral pathways of neuronal signal processing. To obtain multifarious insight across numerous patho physiological process we employed cutting edge approaches such as structural magnetic resonance imaging (MRI): diffusion tensor imaging (DTI), functional MRI (fMRI) and vascular MRI (cerebrovascular reactivity). From the study we obtained several unexpected anatomical and physiological neuro-protective features in the older cohort. Overall, we observed that older cohort exhibit better circum-cerebral structural connectivity as (i) cuneus to superior frontal gyrus (ii) superior frontal gyrus to inferior frontal gyrus (iii) inferior temporal gyrus to inferior frontal gyrus and (iv) cuneus to inferior temporal gyrus. Another, significant neuroprotective anatomical signature was revealed, namely a connectivity through vertical occipital fasciculus across cuneus to temporal cortex in male and female older cohort. Additionally, in older female we observed a pronounced increment in overall functional connectivity network observed in inferior frontal gyrus (both hemispheres). Whereas in older male a substantial increment of functional connectivity network in cuneus was observed as compared to their younger counterpart.

## 1. INTRODUCTION

Ageing accounts for numerous chronic disease (Alzheimer’s disease, stroke, cardiovascular disorders), oftentimes characterized by presence of comorbidities (1,2,3). On average an accelerated brain ageing is reported in female compared to male during adolescence. At the onset of teenage a stark positive age gap is visible in female brain whereas contemporary male brain exhibits a negative age gap (4). Advanced brain age is correlated with the propensity of number of neurodegenerative disease (i.e alzheimer’s) and psychiatric disease (i.e schizophrenia) (5,6,7). Additionally, women tend to accumulate more age-related neurodegenerative disease than men (8). Resting state fMRI analysis also demonstrate considerably different brain regions are functionally correlated with ageing progression in different genders (9,10).

The information processing in brain is mediated through two major parallel streams of neural signal processing the dorsal and ventral pathways. This two-stream hypothesis is most relevant in analysing visual as well as auditory processing. According to the Mishkin-Ungerleider hypothesis this model highlights two major aspects (i) features help in object identification (“What”) processed in occipito-temporal or ventral stream; (ii) perception of “spatial” information are processed in the occipito-parietal or dorsal stream (11). It may be mentioned that further Milner-Goodale (MG) model underscored that the output or task goal itself select the activation of ventral Pathway (Visuo-motor model) (12,13). Dorsal visual stream dysfunction is often associated with impaired complex visual perceptions, diminished visual field (14). These impairments are coupled with several neurodevelopmental (autism spectrum disorders) and neurodegenerative disorders (Alzheimer’s disease) (15). Likewise, impairment in ventral stream is correlated with perusal and visual memory (16).

Notably, human brain undertakes parallel transmission pathways to improve the transmission fidelity and to offset the neurodegenerative features (17). But the underlying mechanism that enable the brain to choose selective pathways over parallel pathways are not well defined. Human brain embroidered a parallel mode of signal processing to interlink somato-sensory and attention regions with executive control and default mode system (18). Additionally, how the brain circuitry coordinate between the selective and parallel pathways of neuronal signal processing prior to perform a function is yet to be deciphered. In our study we try to address how brain structurally and functionally counterbalance the degenerative measures over an age gradient in both genders during the course of normal ageing.

Graph theory is a statistical framework to represent the functional and structural neuronal connectivity in a topological space. It comprises an intricate network that consist of nodes (neurones and brain regions) and connected by edges (synapses or axonal projection). Implementation of graph theory to decode the changes of local and global connectivity throughout the dorsal-ventral pathway may be needed to highlight the key neuroprotective features conserved across the vital cerebral nodal hubs. Our study has demonstrated the preservation of functional connectivity strength in inferior frontal gyrus for older male and in cuneus for older female cohort, over the normal ageing process.

The propensity of neurodegenerative and neurovascular impairment increases with ageing and significantly influence the onset of cognitive deficit like dementia. Intravascular CO2 mediated alteration of cerebral blood flow response under normal physiological condition is termed as cerebrovascular reactivity (CVR). Studies have demonstrated, over the normal ageing process there is a stiffening of central vessels that eventually diminish the cerebrovascular reactivity and consequently, promote microvascular damage (19,20). It is crucial to know how age-wise alteration of CVR may accelerate the commencement of cerebrovascular disease pathophysiology and whether gender differences impart any significant changes in CVR.

We have extensively analysed the and figured out several neuro-restorative signatures across the circumcerebral pathways in elderly population distinct from younger cohort. In this study we made an effort to unravel pronounced neuroanatomical divergence across dorsal-ventral pathways through a wide ageing gradient. These pathways and features that preserve neurorestorative features offer vital insights aimed at promoting healthy cognitive ageing.

## 2. Materials and Methods

### 2.1 Image Acquisition

MRI scans for the particular study were obtained from the Max Plank Institute Mind-Brain-Body database. All the subjects were assessed for standardized clinical interview for DSM IV(SCID-I), Hamilton depression scale and Borderline symptoms list to determine Psychiatric symptoms. The list also consists of 6 cognitive tests as well as 21 questionaries related to emotional behaviour, personality test, eating behaviour and addictive behaviour.

### 2.2 Structural Imaging

High angular resolution whole brain Diffusion weighted images (DWI) form 60 subjects [all healthy, 15 healthy old (age group: 60-70) 15 healthy young (age group: 20-30)] in both male and female cohort, were procured with the help of a 3.0-T magnetic resonance imaging (MRI) scanner (MAGNETOM Verio, Simens Healthcare) equipped with a 32-channel head coil array.

In order to generate structural connectivity, the DWI scans were pre-processed with DSI-studio platform. Reconstruction of diffusion data was performed using generalized q-sampling imaging (GQI) with an angular threshold 60°, step size 0.1 mm, minimum length = 5 mm, maximum length = 300 mm. Afterwards, statistical estimation was performed employing parametric test and Welch’s unpaired t-test for the groupwise comparison of DTI metrices i.e axial diffusivity, number of tracts, volume (mm^3^) between younger and older group.

### 2.3 Functional Imaging

Resting-state functional-MRI study was performed on same 60 participants. Functional connectivity study was undertaken on T1 weighted echoplanar multiband BOLD images. All the participants were in eye-opened awake state looking at a low contrast fixation cross. Sequence parameters were TR=1400ms, Number of volumes:657, Total acquisition time: 15 minutes and 30 seconds. Pre-Processing of brain scan and statistical analysis were conducted using the CONN toolbox, version 22.a (Nieto-Castanon and Whitfield-Gabrieli, 2022) and SPM version 12 (Welcome Department of Imaging Neuroscience, UCL, UK) in MATLAB R2022a (The MathWorks Inc., Natick, MA, United Kingdom).

The entire run consists of several pipelines that include re-alignment and unwrapping for co-registration of all the scans to the first scan. Slice time correction was performed in order to reconcile the time differences of different slices. This is subsequently followed by outlier detection with a conservative setting (95th percentiles), segmentation (white matter, gray matter, and cerebral spinal fluid), normalization according to the Montreal Neurological Institute atlas (MNI 152) and smoothing at 8 mm full width half maximum (FWHM) Gaussian kernel to diminish the noise from each voxel’s time series signal. After the preprocessing steps, default blister variables of CONN toolbox were employed to perform denoising here, each signal is then filtered through a 0.01 Hz band-pass filter.

We compared several graph theory metrices of different older and younger groups in both male and female cohort in order to estimate the functional connectivity strength. The metrices are global efficiency, local efficiency, eigenvector centrality, closeness centrality, betweenness centrality. The adjacency matrix threshold is set as 0.15 cost to initially threshold the edges between nodes, whereas the analysis threshold is set as p<0.05. Then we employed parametric test and welch’s unpaired t-test for the groupwise comparison for statistical estimation.

### 2.4 Cerebrovascular Reactivity

Vascular reactivity is performed in FMRIB software library (Oxford University, UK). Anatomical images (T1 images) of all the subjects are processed through Brain Extraction Tool (BET) at a fractional intensity threshold of 0.5 in order to remove non-brain tissue from whole head. Then using hidden Markov random field model, the T1_bet image is segmented into three different tissue type (Grey Matter, White Matter, CSF) using partial volume image. Furthermore, we employed FEAT (FMRI expert analysis tool) interface to perform pre-processing at a spatial smoothing FWHM of 5mm and registration. Hence, we treated CSF signal as stimulus signal to perform our vascular reactivity analysis. Using the CSF stimulus we have created a binary mask for CSF, grey matter and white matter. The study is followed by calculation of mean CSF values, mean GM and WM value of each volume. We adjust the Z-Threshold Value to 2.3 to create a brain activation map with cerebrospinal fluid acting as the explanatory variable.

## 3. RESULTS

### 3.1 Functional Connectivity

Based on ROI-ROI analysis in the male and female cohorts we have obtained significant alteration of functional connectivity in four nodal points of dorsal streams (cuneus, superior frontal gyrus) and ventral streams (inferior Temporal gyrus, inferior Frontal Gyrus) in both hemispheres that is evident from table1,table2,table3 and table4.

**Table. 1.** Functional connectivity measures of Dorsal and Ventral nodal points in elderly male brain.

| Brain Regions | Beta value | T(14) | p-unc | p-FDR |
| --- | --- | --- | --- | --- |
| Cuneus (right) | 0.36 | 9.47 | 0.0000 | 0.000003 |
| Cuneus(left) | 0.35 | 7.98 | 0.000001 | 0.000016 |
| Superior Frontal Gyrus(right) | 0.13 | 4.14 | 0.000998 | 0.003037 |
| Superior Frontal Gyrus(left) | 0.13 | 3.34 | 0.004819 | 0.011428 |
| Inferior Frontal Gyrus(right) | -0.12 | -4.82 | 0.000273 | 0.001079 |
| Inferior Frontal Gyrus(left) | -0.13 | -5.14 | 0.000151 | 0.000661 |
| Inferior Temporal Gyrus(right) | 0.11 | 4.54 | 0.000466 | 0.001682 |
| Inferior Temporal Gyrus(left) | 0.15 | 5.23 | 0.000127 | 0.001682 |

**Table. 2.**
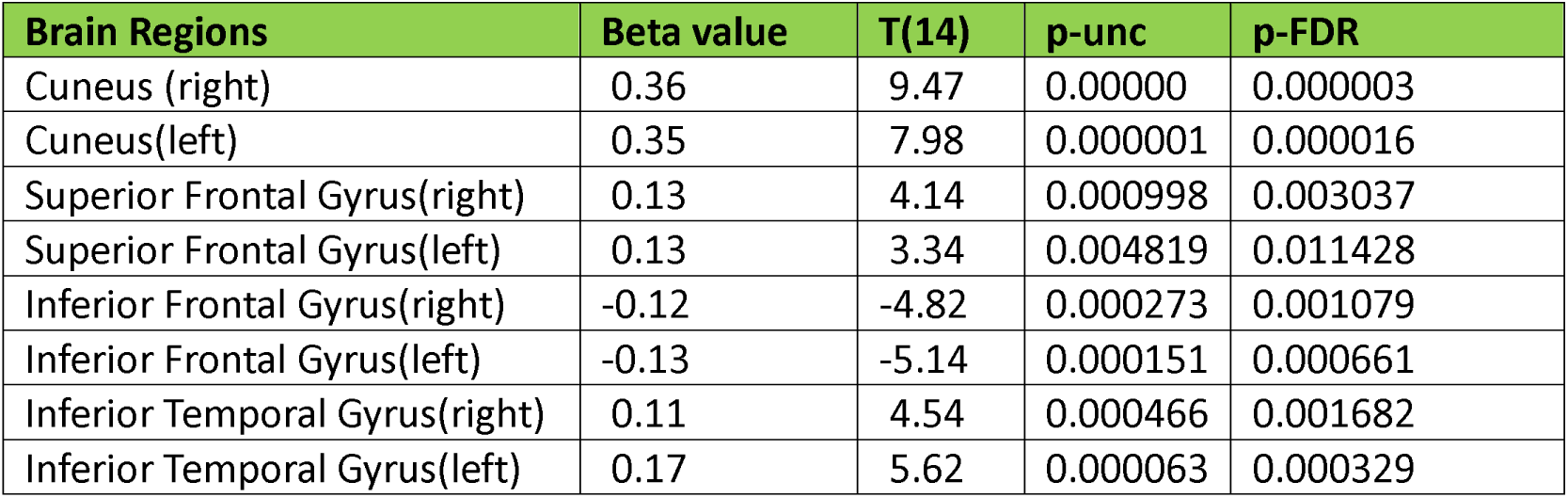
Functional connectivity measures of Dorsal-Ventral nodal points in elderly female brain.

| Brain Regions | Beta value | T(14) | p-unc | p-FDR |
| --- | --- | --- | --- | --- |
| Cuneus (right) | 0.36 | 9.47 | 0.00000 | 0.000003 |
| Cuneus(left) | 0.35 | 7.98 | 0.000001 | 0.000016 |
| Superior Frontal Gyrus(right) | 0.13 | 4.14 | 0.000998 | 0.003037 |
| Superior Frontal Gyrus(left) | 0.13 | 3.34 | 0.004819 | 0.011428 |
| Inferior Frontal Gyrus(right) | -0.12 | -4.82 | 0.000273 | 0.001079 |
| Inferior Frontal Gyrus(left) | -0.13 | -5.14 | 0.000151 | 0.000661 |
| Inferior Temporal Gyrus(right) | 0.11 | 4.54 | 0.000466 | 0.001682 |
| Inferior Temporal Gyrus(left) | 0.17 | 5.62 | 0.000063 | 0.000329 |

**Table. 3.**
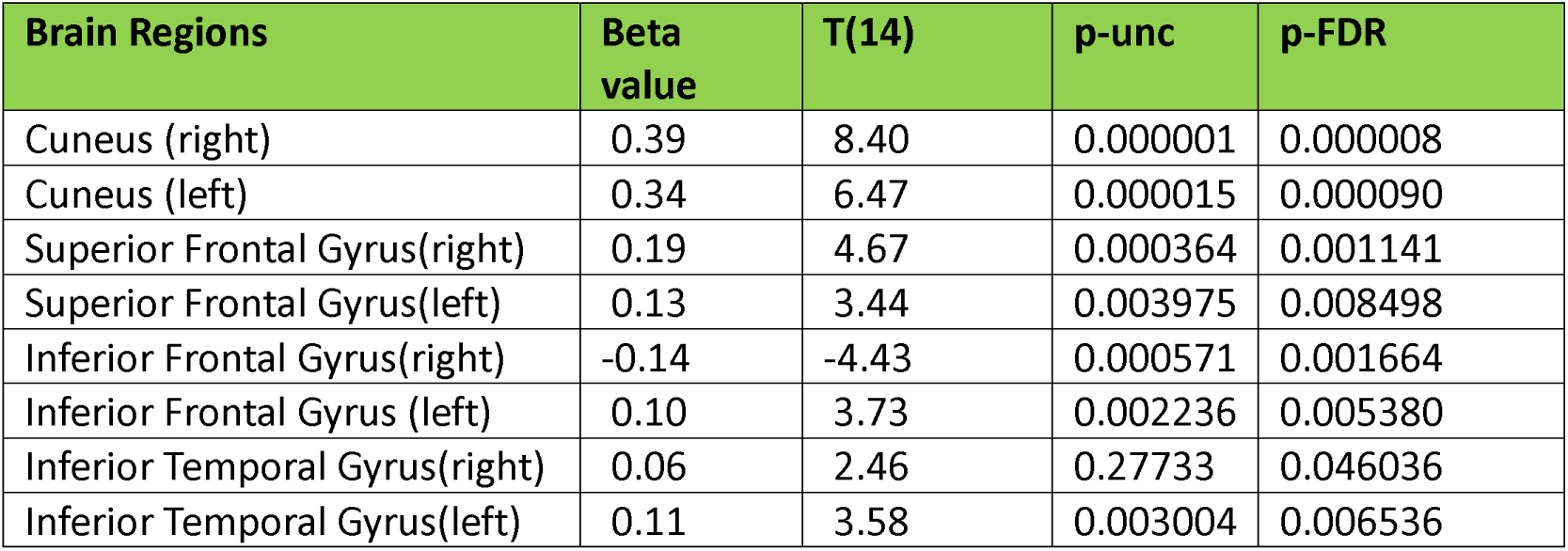
Functional Connectivity measures of Dorsal-ventral nodal points in younger male brain.

**Table. 4.** Functional connectivity measures of Dorsal-Ventral nodal points in young female brain.

| Brain Regions | Beta value | T(14) | p-unc | p-FDR |
| --- | --- | --- | --- | --- |
| Cuneus (right) | 0.32 | 8.85 | 0.00000 | 0.000006 |
| Cuneus(Left) | 0.29 | 6.62 | 0.000012 | 0.000101 |
| Superior Frontal Gyrus(Right) | 0.09 | 2.51 | 0.024851 | 0.052888 |
| Superior Frontal Gyrus(Left) | 0.05 | 1.10 | 0.290569 | 0.401954 |
| Inferior Frontal Gyrus(Right) | -0.15 | -3.87 | 0.001710 | 0.005792 |
| Inferior Frontal Gyrus(left) | -0.13 | -3.54 | 0.003295 | 0.009272 |
| Inferior Temporal Gyrus(Right) | 0.11 | 3.09 | 0.008026 | 0.020187 |
| Inferior Temporal Gyrus(Left) | 0.11 | 2.34 | 0.30392 | 0.070291 |

Age-wise functional activation in focal brain regions is as follows (Fig1).and Supplement1, Supplement2, Supplement3, Supplement4

**Fig. 1.**
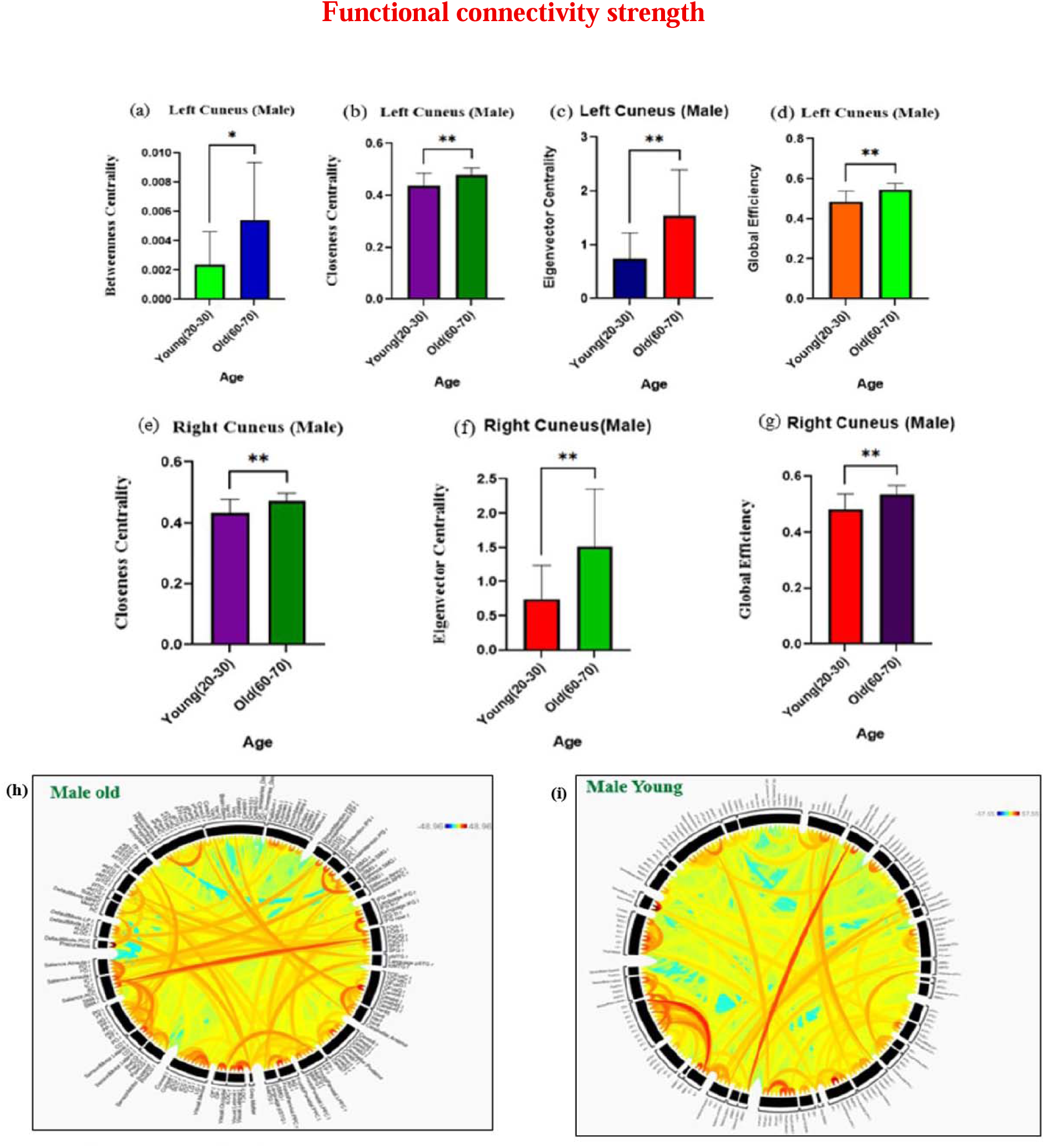
Increased in functional connectivity of cuneus in old age males: ***Left cuneus***:(a) betweenness centrality index;(b) closeness centrality index;(c) eigenvector centrality index;(d) Global efficiency index. ***Right cuneus***: (e) closeness centrality; (f) eigenvector centrality;(g) global efficiency. ***Connectogram***:(h) Old male (i) Young male. It is also noted that the number of very long connectivity fibers in older males are more divergently distributed than in younger males, indicating increased long collateral connections in older males.

#### 3.1.a. Variation in Intra and Inter nodal functional connectivity in female

We employed graph theory parameters to dissect the strength of intra-inter network functional connectivity across young and old female cohort. We found that in older in older female there is a substantial increment in salience network connectivity in both hemispheres. Further in this subject group in older females there is an escalated minuscule affinity (Betweenness centrality index) in Left inferior frontal gyrus (pars opercularis) as compared to younger females. Additionally, there is an enhancement kinship for vital nodes (eigenvector index) with right inferior frontal gyrus. Moreover, there is a surge in extensive integration (global efficiency index) of this region of interest (Right inferior gyrus) connected with overall brain (Fig.2).

**Fig. 2.**
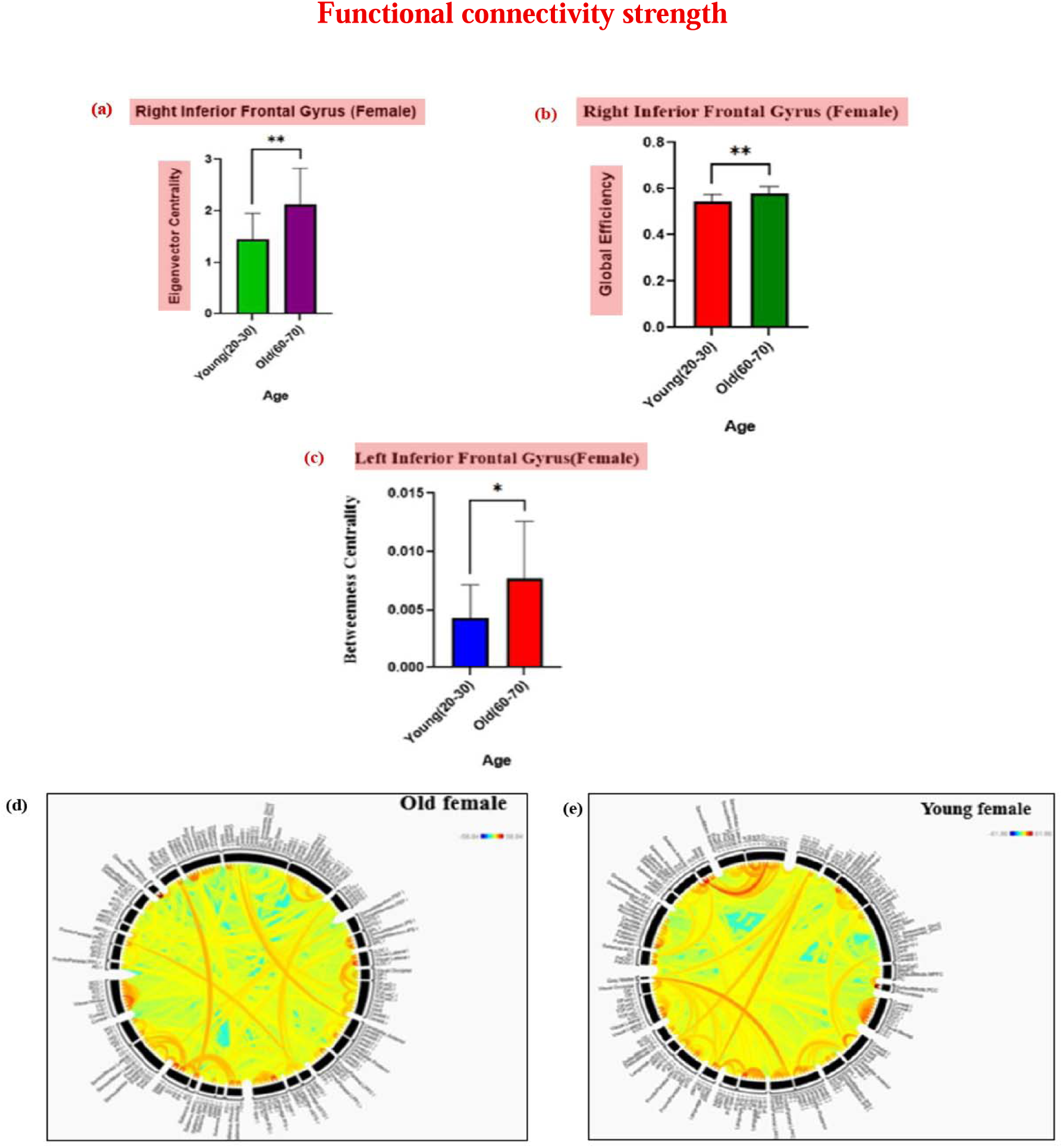
Increase in functional connectivity of inferior frontal gyrus in old age females: ***Right inferior temporal gyrus:***(a) eigenvector centrality index; (b)global efficiency index. Left inferior temporal gyrus: (c) betweenness centrality index. ***Connectogram***:(h) Old female (i) Young female. Furthermore, it may be mentioned that the number of very long connectivity fibers in older females are more divergently distributed than in younger females, indicating increased long collateral connections in older female.

#### 3.1.b. Variation in Intra and Inter nodal functional connectivity in male

A noteworthy increment of graph theory based functional connectivity default mode network (DMN) precisely in cuneus is observed in older male cohort compared to the younger group.

A prominent surge in minuscule affinity (Betweenness centrality) was discerned in left cuneus whereas an increment in other parameters like eigenvector index, global efficiency index and closeness centrality index was noticed in both hemisphere of default mode network (Cuneus) of older cohort in as compared to younger one (Fig.1).

### 3.2 Structural Alteration

#### 3.2.a. Dorsal stream (Anterior-Posterior segment)

The segment that connects cuneus in the posterior cerebrum and superior frontal gyrus in anterior cerebrum exerts higher Axial diffusivity in older female group as compared to the younger one (p< 0.04), account for an improved axonal integrity in older cohort. Besides, an increment in axial diffusivity was also observed in the same bridging tract was also observed within older male cohort (p<0.008)(Fig:3a,3b).

#### 3.2.b. Intra-frontal connectivity

A number of diffusion-metrices synonymous with neuroprotection were analyzed through intra-frontal connectivity that connects superior frontal gyrus to pars opercularis of Inferior frontal gyrus. Notably, older female cohort have significantly higher axial diffusivity in left (p<0.0077) hemisphere (Fig:5b). Furthermore, tract count in the left hemisphere were significantly greater (P< 0.02) in older subject, that accentuated a strive towards age-wise neuro-compensatory mechanism (Fig:5a).

#### 3.2.c. Ventral stream (Fronto-temporal segment)

The segment connecting inferior temporal gyrus in temporal lobe to the pars opercularis of inferior frontal gyrus in the frontal lobe demonstrated a significant increment of axial diffusivity (p< 0.02) in older female cohort in right hemisphere as compared to their younger counterpart(Fig:3d). This disjunct in axial diffusivity is an indicator of revamped neuronal integrity, potentially an age-related adaptation to offset white matter structural perturbation.

#### 3.2.d. Occipitotemporal connectivity

Nerve tracts in the left hemisphere linking cuneus in the occipital pole and inferior temporal gyrus in temporal lobe display a significant increment in tract volume in older cohort (p< 0.009) compared to the younger cohort(Fig:3c). This anatomical reinforcement with the age may signify an age wise reorganization of neuronal connectivity for better cognitive and functional outcome in elder individuals. Another noteworthy finding of our study is highlighted by the abundance of vertical occipital fasciculus, a vital fibre tract involved in complex visual processing in older subjects of both male and female cohort. Whereas the nerve fibre is absent in younger population (Fig.4).

**Fig. 3.**
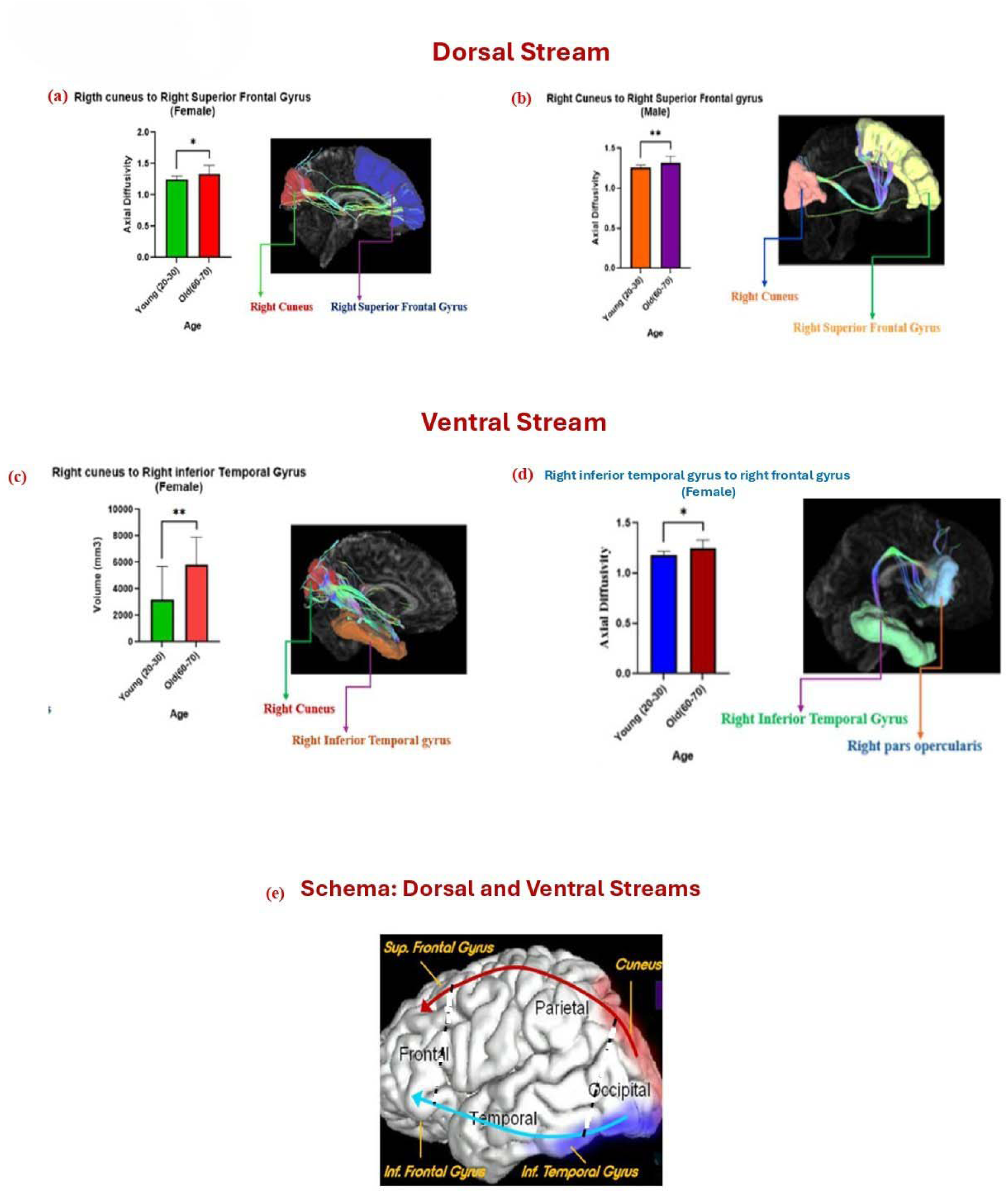
Increased in structural connectivity of dorsal stream and in ventral stream in old age. ***Dorsal stram:*** (a) right cuneus to right superior frontal gyrus in females;(b) right cuneus to right superior frontal gyrus in males; ***Ventral stream:*** (c) posterior segment of ventral stream : right cuneus to right inferior temporal gyrus in females; (d)Anterior segment of ventral stream: right inferior temporal gyrus to right inferior frontal gyrus (pars opercularis); (e) Schema of dorsal and ventral sreams across the brain.

**Fig. 4.**
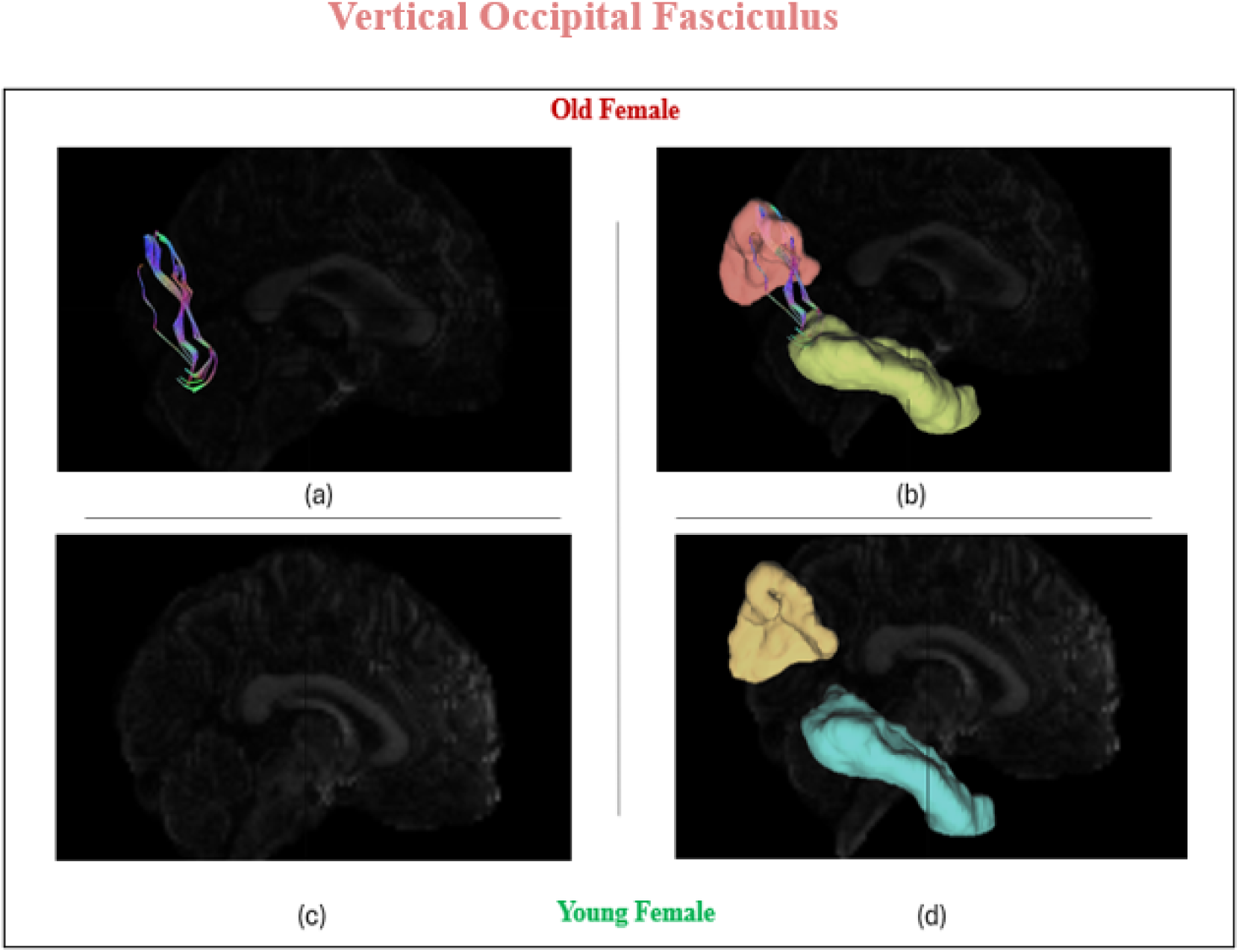
Preservation of **<u>vertical occipital fasciculus</u>** across elderly cohort (male and female) (a) Vertical occipital fasciculus of ventral stream. (b) ***Vertical Occipital fasciculus*** :connecting cuneus and inferior temporal gyrus. (c) Young female brain dwi scan. (d) Vertical occipital fasciculus is absent in young female cohort.

### 3.3 Cerebro-vascular Reactivity

In both the cohort no significant alteration of age-wise vascular reactivity was observed (Male CVR: old vs young p<0.1; Female CVR: old vs young p<0.09). A comparison between both genders demonstrated preservation in perfusion during the course of ageing. We noticed no significant changes of gary matter CVR in male young vs old p<0.4 and white matter p<0.2. Similarly, no significant changes of gray matter CVR in female young vs old p<0.2 and white matter p<0.5 (Fig6, Fig7).

**Fig. 5.**
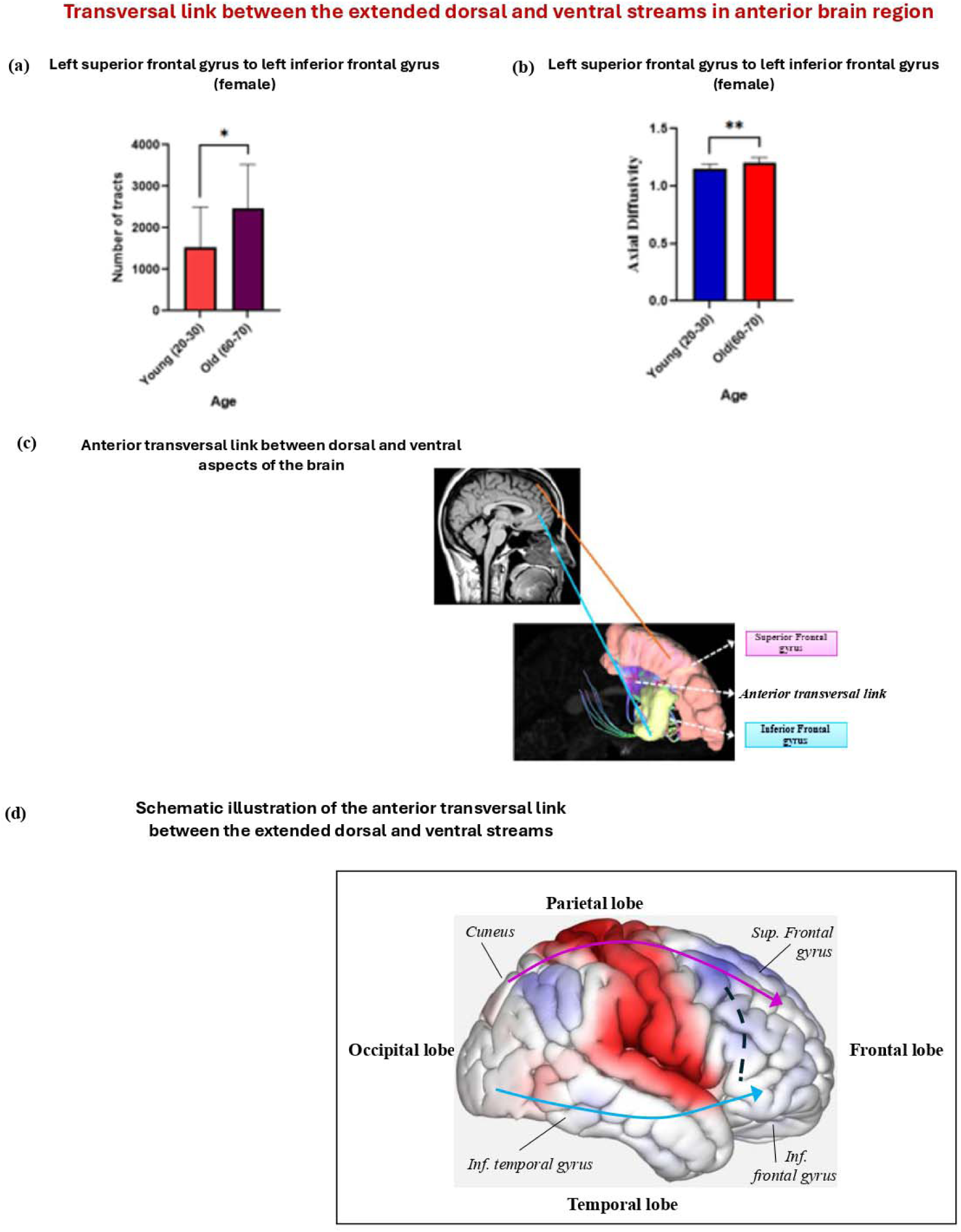
Increase in structural connectivity of th anterior transversal link between the extended dorsal and ventral streams in old age: (a) anterior transversal link between left superior frontal gyrus to left inferior frontal gyrus (opercular region).The number of tracts is estimated for the anterior link. (b) Anterior transversal link between left superior frontal gyrus to left inferior frontal gyrus(opercular region). The axonal integrity is estimated for the anterior link. (c) Tractographic representation of anterior transversal link. (d) Schematic illustration of the anterior traversal link(dashed line) between the extended dorsal and ventral streams.

**Figure 6:**
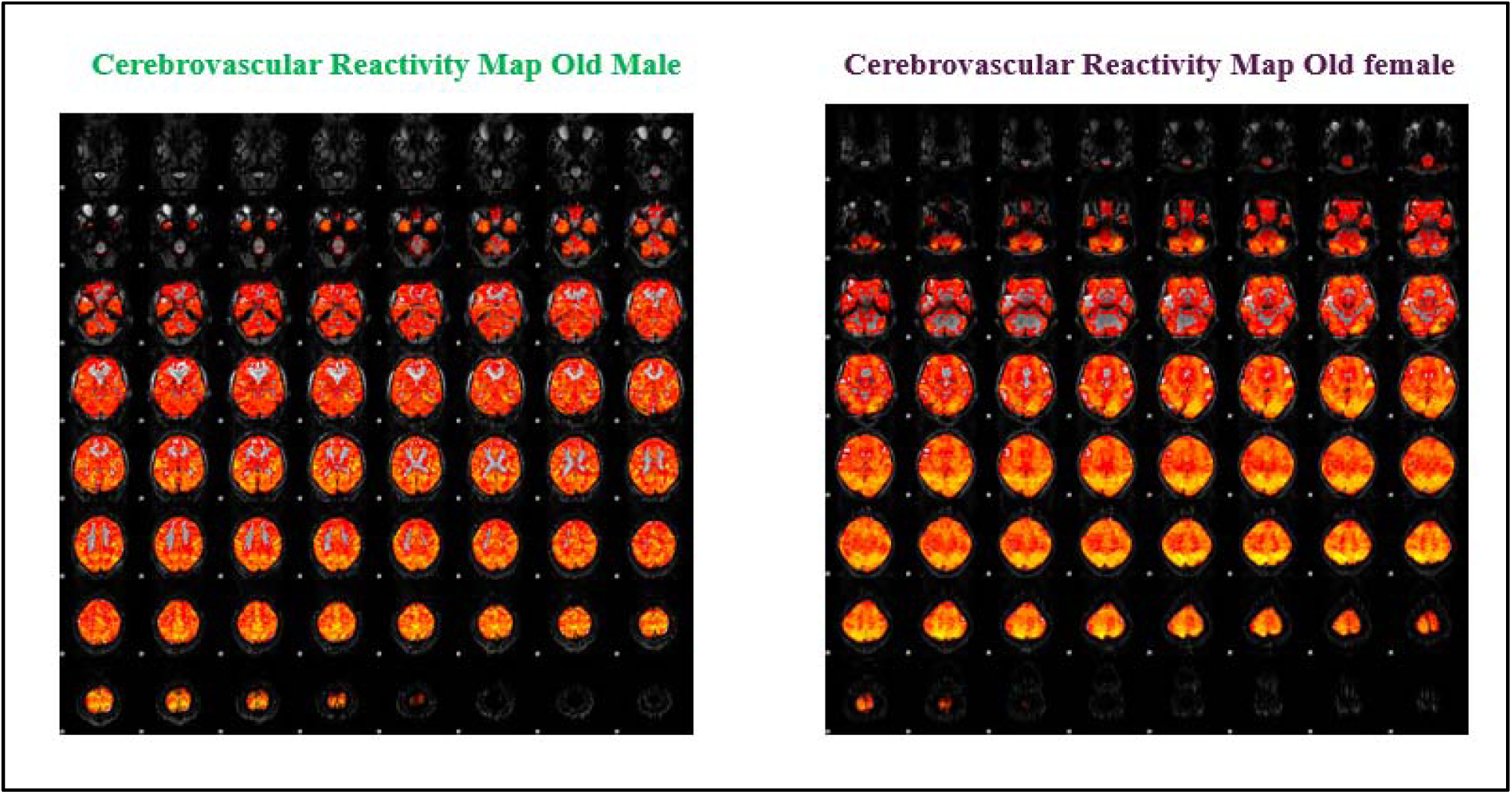
Cerebrovascular Reactivity map in old cohort (both male and female)

**Figure 7:**
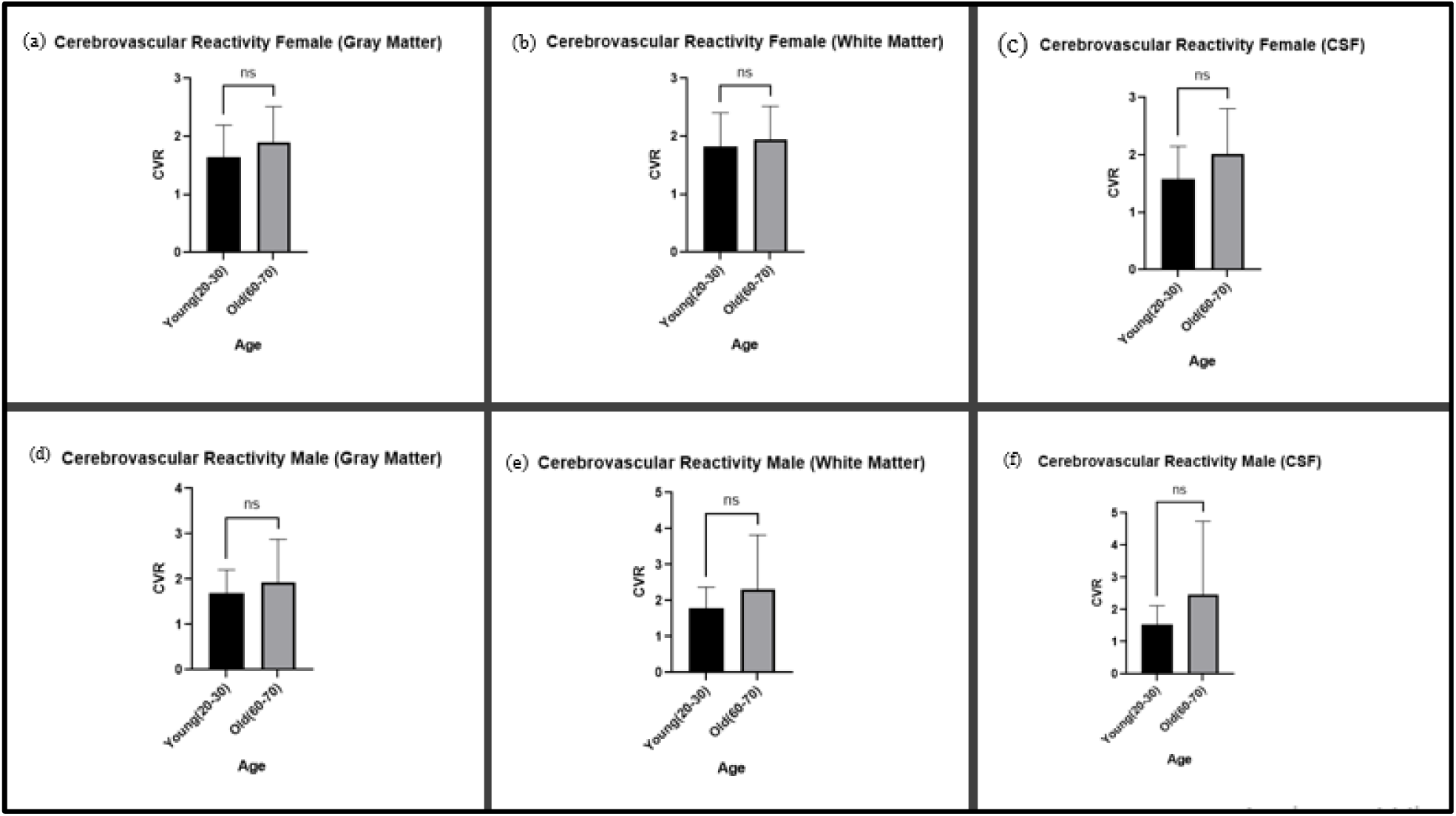
Cerebrovascular reactivity no significant changes in (a) female grey matter, (b) white matter, (c) cerebrospinal fluid. Similarly, we didn’t obtain any changes in male older cohort as compared to the younger in (d) grey matter, (e) white matter and (f) cerebrospinal fluid.

## 4. DISCUSSION

In our study we have demarcated a significant increment of functional connectivity within salience network precisely in Inferior Frontal gyrus of both hemisphere in elder female cohort as compared to younger female. Right Inferior Frontal gyrus is a vital node that facilitate inhibitory control, an ability to control already initiated action (21). These improved connectivity in elderly may account for adaptive tactics of the old brain to preserve its executive function. Patients affected with bipolar disorder exhibited an impaired functional connectivity of IFG with in the fronto limbic and fronto-striatal network (22), Our study has highlighted IFG exerted an improvement in miniscule connectivity, Global and vital connectivity an attainment to ameliorate the cognitive pliancy.

The default mode network (DMN), a fundamental neurobiological system or a baseline network active in the brain at resting state, the network constructs the internal narrative of ‘selfdom’ (23). exhibits a significant change during ageing in male older cohort. A marked increment of betweenness centrality in both hemispheres of cuneus signify a pivotal role of cuneus in integrating information across different brain network. The observation corroborates the impression that during the course of ageing old brain depends heavily on specific nodes to maintain cognitive performances potentially compensating for age-wise neurodegeneration in other brain areas (24,25,26).

Besides, enhancements in global efficiency and eigenvector centrality in both hemispheres of the cuneus further align with the notion that old brains undergo a reconstitution of functional brain networks to sustain cognitive performance (27).

In both dorsal and ventral streams, we have noticed a significant increment of Axial diffusivity in older cohort both in male and female. Axial diffusivity is a parameter of neuronal integrity. It was reported that white matter tracts with higher axial diffusivity facilitates the transition of slow wave neuronal signals into upstate (28). Whereas, in mice model reduction of axial diffusivity is correlated with neuronal injury (29,30). An important study highlighted, the reduction of axonal fiber tortuosity may give rise to more straightened neuronal fiber and subsequently increase the axial diffusivity with the progression of ageing. On the contrary, in the early age the event of axonal pruning and cytoskeletal development may result in diminished axial diffusivity (31,32,33). Increment of nerve tract volume spanning through left Inferior frontal Gyrus (pars opercularis) to Left Inferior temporal gyrus of ventral stream in older female cohort is a culminative neurorestorative measure adopted by brain against hyperosmolarity or hypernatremia (34).

The vertical occipital fasciculus is major nerve tract that link the vital nodal points of dorsal(cuneus) and ventral stream (Inferior Temporal Gyrus). Yeatman for the first time highlighted the presence VOF and its anatomical alignment in living human brain (35). VOF intersect the visual word from area (VWFA) of occipital temporal sulcus involved in skilled reading (36). Whereas, Greenblatt put the first light on the physiological significance of VOF, he discerned that patient with a brain tumor distorted its VOF and patient with surgical resection of VOF succumb to pure alexia (37,38).

Cerebrovascular reactivity (CVR) is a measure to the elasticity of the cerebral vessels under the influence of blood CO2 partial pressure or vasoactive substances. It is vital to maintain normal respiration and to shorten the periods of nonbreathing. In mice impaired cerebrovascular reactivity account for abnormal breathing and respiratory dysfunction. In human the changes in vascular dynamics impact speaking, sighing and subsequently the pH in brain extracellular fluid (39). In our study we have observed no significant difference in cerebrovascular reactivity in older cohort as compared to the younger one in both genders. This conservation in vascular reactivity may help to diminishing chemo-sensitive fear and anxiety like behaviour in-order to balance shift in blood CO2/pH levels (40).

In conclusion, this manuscript contributes significantly to ageing dynamics of brain coupled with neuroprotection by elucidating the complex interplay between structural and functional connectivity in older adults. The findings underscore the importance of considering integrity of dorsal and ventral stream of neural pathway in aging research and highlight the potential for neuroimaging to inform targeted interventions. As we continue to unravel the mysteries of the aging brain, it is imperative to adopt a multifaceted approach that encompasses both biological and psychosocial factors, paving the way for innovative strategies to enhance cognitive health in the aging population.

## SUPPLEMENTARY MATERIALS

**Figure supplement.1.**
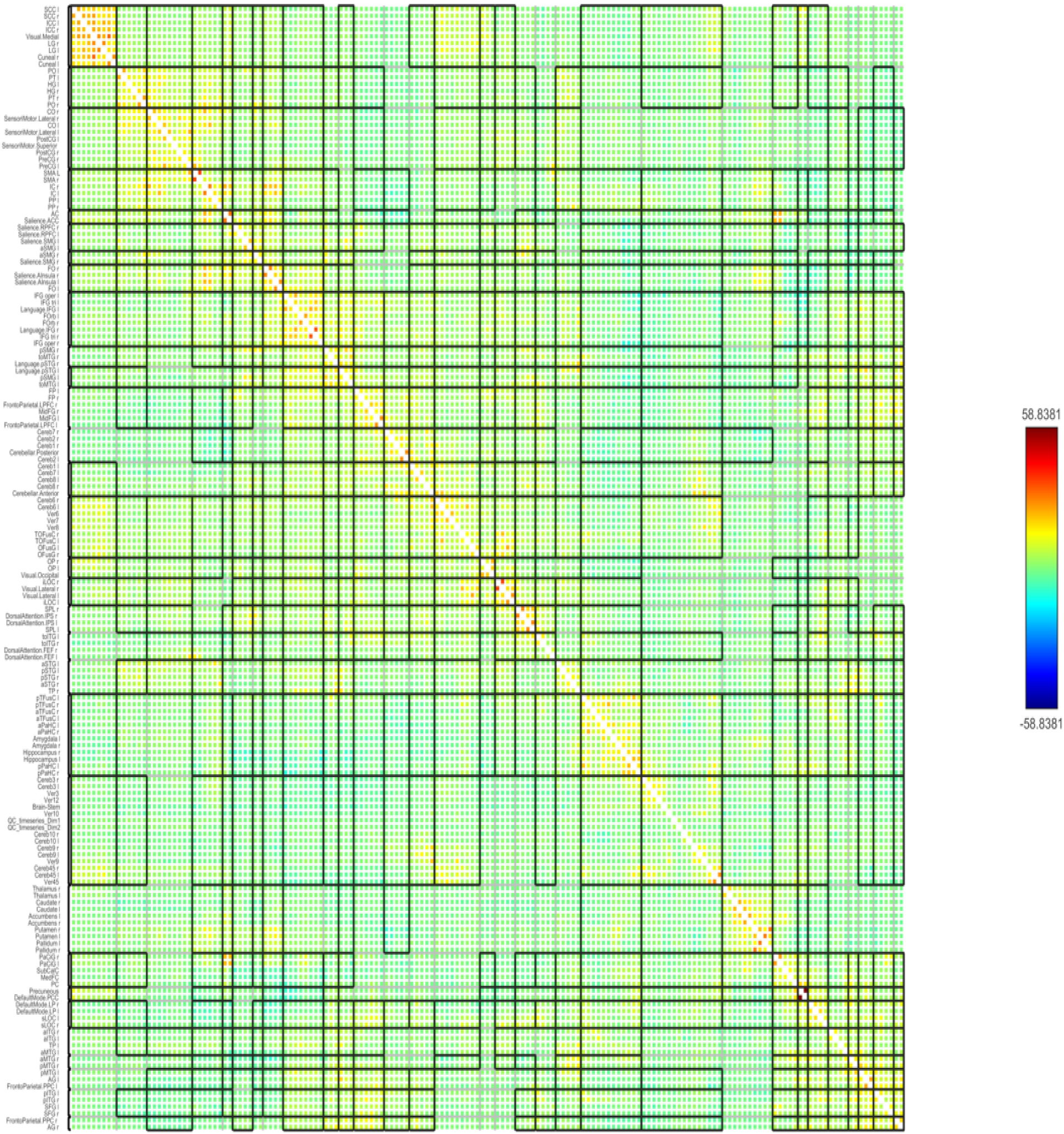
ROI to ROI functional connectivity matrix in old female (60-70 years) cohort, with advanced Family wise Error control setting, Cluster threshold p< 0.05, two-sided, p-FDR corrected.

**Figure supplement.2.**
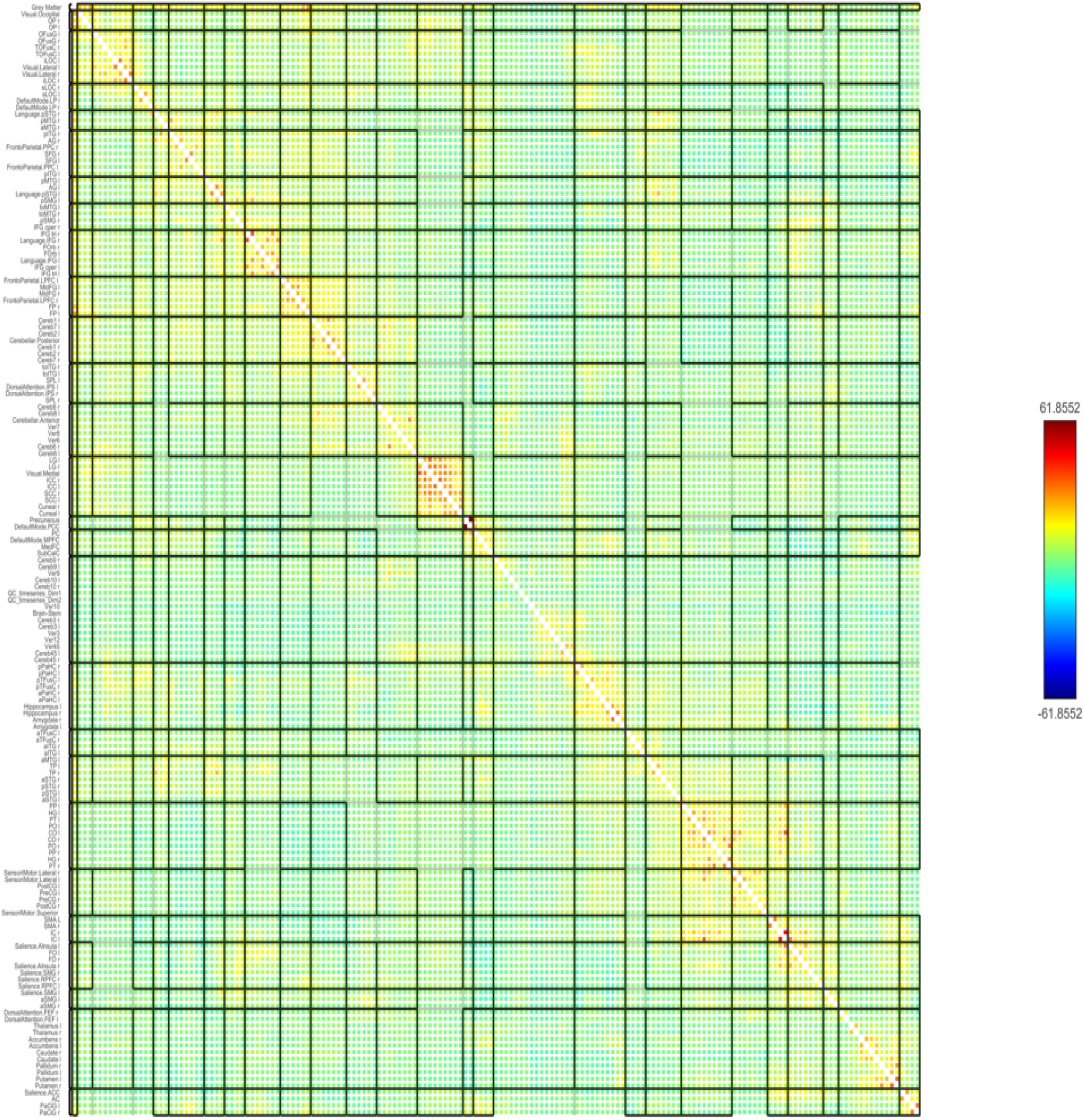
ROI to ROI functional connectivity matrix in young female (20-30 years) cohort, with advanced Family wise Error control setting, Cluster threshold p< 0.05, two-sided, p-FDR corrected.

**Figure supplement.3.**
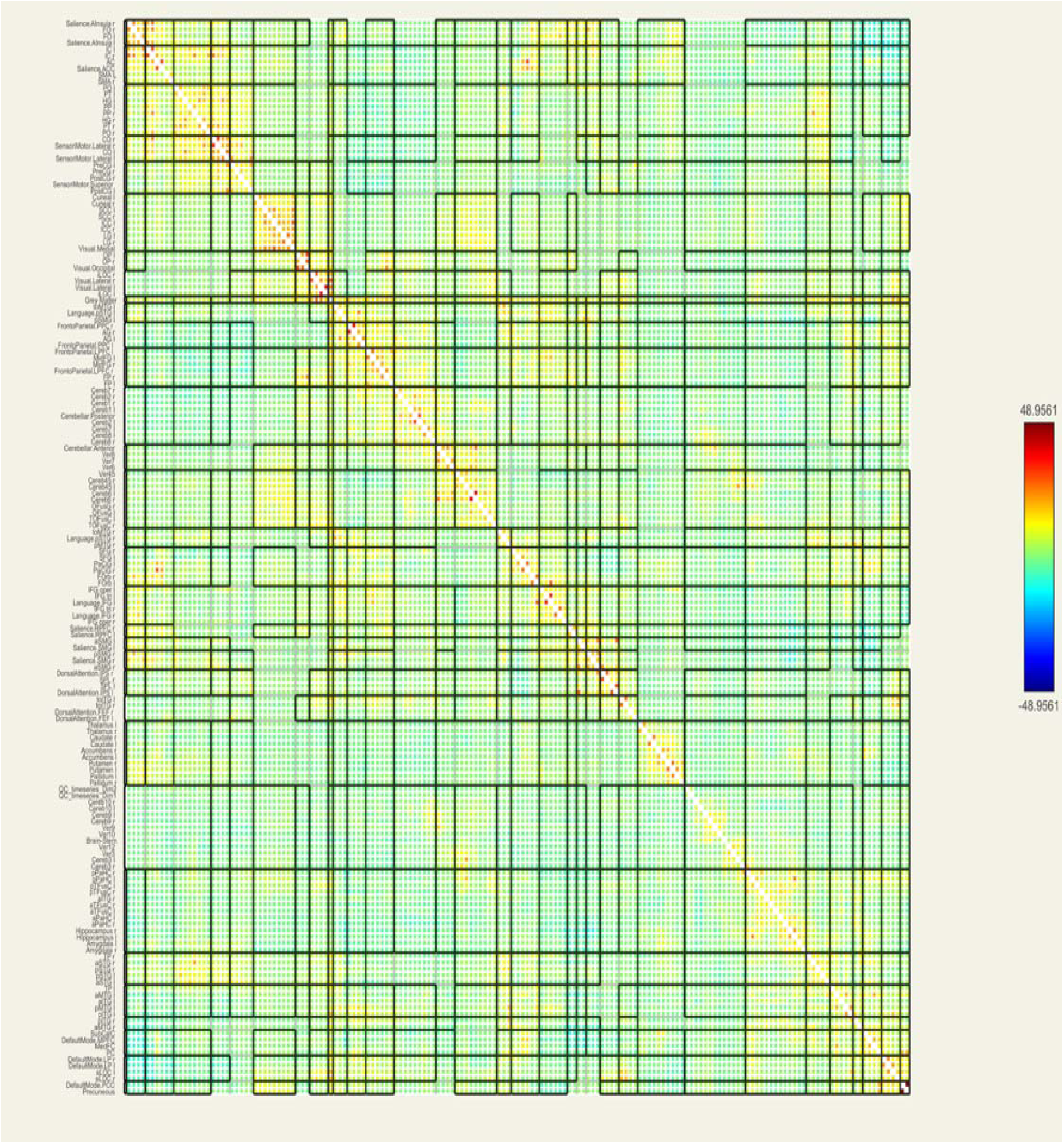
ROI to ROI functional connectivity matrix in Old Male (60-70 years) cohort, with advanced Family wise Error control setting, Cluster threshold p< 0.05, two-sided, p-FDR corrected.

**Figure supplement.4.**
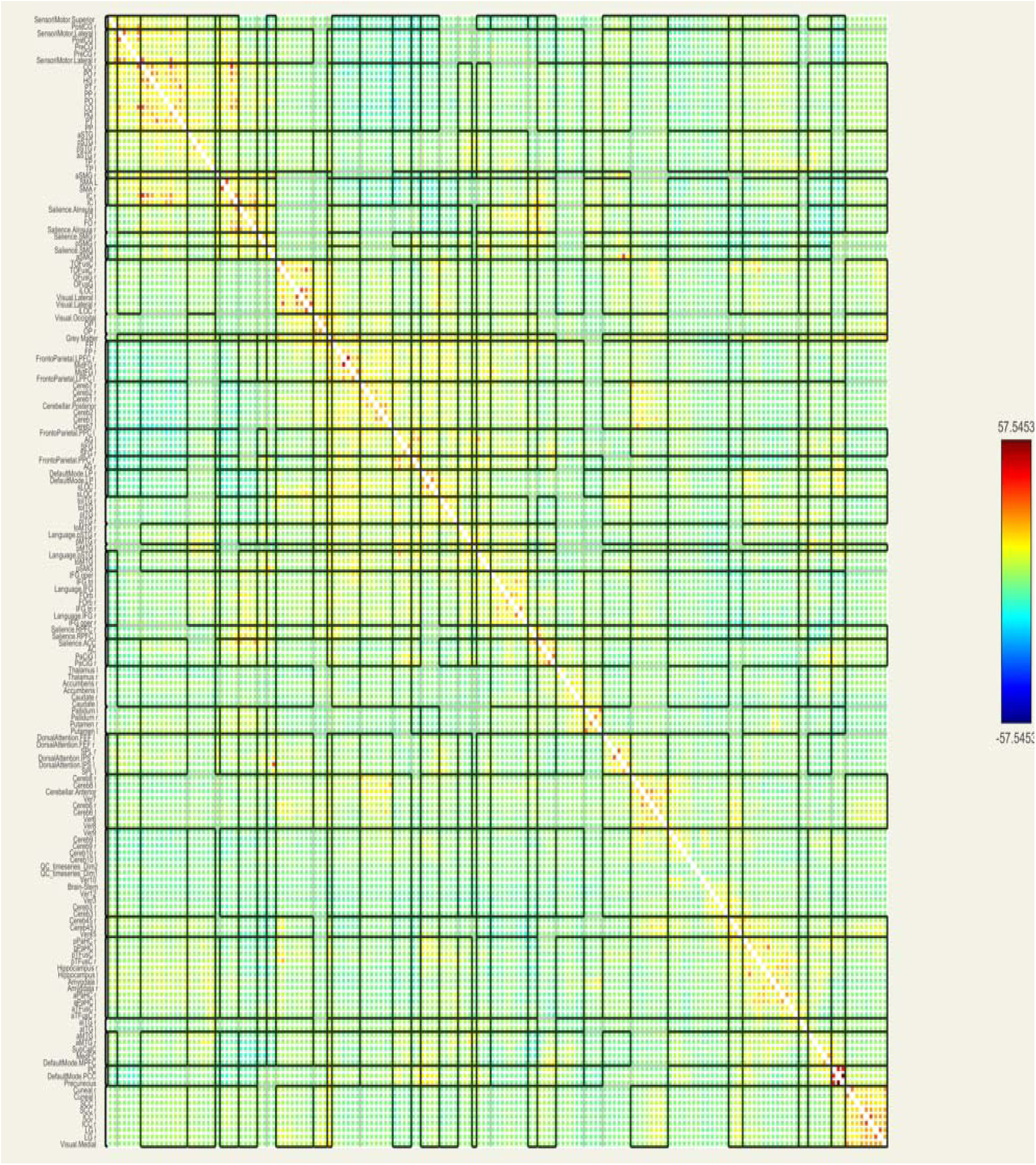
ROI to ROI functional connectivity matrix in Young Male (20-30 years) cohort, with advanced Family wise Error control setting, Cluster threshold p< 0.05, two-sided, p-FDR corrected.

